# The genetics of behavioral isolation in an island system

**DOI:** 10.1101/250852

**Authors:** Thomas Blankers, Kevin P. Oh, Kerry L. Shaw

**Affiliations:** Department of Neurobiology and Behavior, Cornell University, Ithaca, NY, USA

**Keywords:** speciation, sexual isolation, quantitative genetics, transcriptome, *Laupala*

## Abstract

Mating behavior divergence can make significant contributions to reproductive isolation and speciation in various biogeographic contexts. However, whether the genetic architecture underlying mating behavior divergence is related to the biogeographic history and the tempo and mode of speciation remains poorly understood. Here, we use quantitative trait locus (QTL) mapping to infer the number, distribution, and effect size of mating song rhythm variation in the crickets *Laupala eukolea* and *L. cerasina*, which occur on different islands (Maui and Hawai’i). We then compare these results with a similar study of an independently evolving species pair that diverged within the same island. Finally, we annotate the *L. cerasina* transcriptome and test whether QTL fall in functionally enriched genomic regions. We document a polygenic architecture behind song rhythm divergence in the inter-island species pair that is remarkably similar to that previously found for an intra-island species pair in the same genus. Importantly, QTL regions were significantly enriched for potential homologs of genes involved in pathways that may be modulating cricket song rhythm. These clusters of loci could constrain the spatial genomic distribution of genetic variation underlying cricket song variation and harbor several candidate genes that merit further study.

## 1. Introduction

Behavioral divergence can produce significant reproductive barriers in animals and can be an important force driving speciation [1–5]. However, the genetic mechanisms leading to behavioral divergence, suppression of interspecific recombination, and ultimately the origin of new species remain poorly understood [3, 6–9]. We can make significant progress in understanding the speciation process by dissecting and comparing the genetic architecture of behavioral isolation – the number, effect size, and genomic distribution of loci controlling mating behavior divergence – across a range of different speciation events and by identifying potentially causal genes.

The genetic architecture of phenotypes can be categorized as Type I (polygenic and additive) or Type II (oligogenic and epistatic) architectures [10]. It has been argued that the type of genetic architecture can be informative about the tempo and mode of speciation [11,12]. All else being equal, an oligogenic basis would allow for rapid phenotypic evolution, because the phenotypic effect of a single mutation is expected to be larger; theoretical models for speciation by sexual selection often assume such simple genetic architectures [13–16]. Moreover, it has been suggested that Type II architectures may be lynch pins in radiations involving founder effects [11], such as those accompanying the colonization of archipelagos, due to the volatile evolutionary effects of epistatic interactions under fluctuating demography. In contrast, empirical studies show that divergent sexual traits, which can be an important cause of rapidly evolving behavioral isolation [4], are typically associated with Type I genetic architectures [3,17], without a disproportionate contribution from sex-linked loci [3,18] albeit, the role of X-effects in the evolution of secondary sexual characters remains somewhat contentious, e.g. [19]. A polygenic architecture would allow for gradual (but not necessarily slow) and orchestrated divergence of male and female sexual traits due to the predictable, biometrical action of genetic variants on diverging traits [20,21].

Other genomic influences may also affect the evolution of reproductive barriers. The number of independent genetic factors and the rate of phenotypic evolution can be strongly affected when causal loci cluster in specific genomic regions. Fisher [22] suggested that genes associated with the gradual adaptive change of complex phenotypes will become physically linked over time. He imagined this linkage would maintain the identity of groups with complex phenotypic differences (e.g. divergent populations, dimorphic sexes) by reducing the disruptive effects of recombination in the region with which the phenotype is associated. Indeed, linkage disequilibrium within tightly linked clusters of quantitative trait loci (QTL) can be maintained by low recombination, for example due to chromosomal inversions [23]. The involvement of gene clusters has been implicated in the rapid evolution of (co)adaptive phenotypes in plants and animals [24], including courtship behavior [25–28]. Such clusters allow for orchestrated adaptive responses by reducing recombination between co-adaptive alleles and may be an important mechanism underlying natural variation in complex traits, such as altruistic signals. i.e. ‘green beard’ phenotypes [29], behavior, and co-evolving sexual signals and preferences [21,22,30].

Here, we examine the genetic basis of sexual trait divergence in the rapidly diversifying, endemic Hawaiian cricket genus *Laupala.* We test whether the biogeographic context of speciation can predict the genetic architecture of courtship song divergence and whether the distribution of causal loci is potentially constrained by clustering of functionally related genes. Differentiation in acoustic sexual signaling behavior appears to play a central role in *Laupala* speciation, a group with one of the highest speciation rates documented for arthropods [31]. Species are endemic to single islands and show striking differentiation in male song and female (song) preference [31,32]. Like most other cricket species, the male produces a song by rubbing together its specialized forewings. The songs of *Laupala* crickets are relatively simple and consist of trains of pulses, each pulse produced by a single closing movement of the wings. Evidence suggests that a critical trait used in female mate choice is the repetition rate of these pulses, i.e. the pulse rate [33,34], which is highly heritable and constitutes one of the major sources of phenotypic divergence among *Laupala* species [35–39].

So far, we have not been able to pin cricket song rhythm variation down to individual genes. However, the neurobiology of cricket song generation is generally well understood, involving contributions from descending brain neurons, specialized motor neurons in the thorax called central pattern generators, and related neuromuscular modulators such as ion channels and synaptic targets [40–44]. Therefore, other organisms with known genetic and neurological pathways that drive sex-specific and (acoustic) courtship behaviors can shed light on the potential mechanisms at play in *Laupala* song divergence. In *Drosophila melanogaster*, sexual dimorphism in the nervous system, driven by the interaction between the transcription factors *doublesex* and *fruitless*, provides a developmental foundation for courtship song [45,46] However, neither of these genes contributes to interspecific differences in song rhythm [47]. Other genes, including *slowpoke* and *cacophony*, have been implicated in species and population differences across *Drosophila* species: these genes control the properties of ion (calcium and potassium) channels at neuromuscular junctions [48–51] and can transform the output of neuronal networks including central pattern generators [52]. Therefore, we may expect similar neuromodulators as well as genes involved in synaptic transmission at neuromuscular junctions and rhythmic behavior to contribute to variation in song rhythm in crickets.

The first goal of this study is to illuminate the genetic architecture of sexual trait divergence in light of different geographic modes of speciation. Despite rapid phenotypic divergence between *L. cerasina* and *L. eukolea* following colonization of a new island [53], evidence from a previous biometric study shows that multiple genetic factors (˜5) underlie pulse rate variation between these species [39]. We build on this knowledge using quantitative trait locus (QTL) mapping to examine the number, effect size, genomic distribution and interactions of loci contributing to interspecific pulse rate variation between *L. cerasina* and *L. eukolea*, in order to characterize the genetic architecture as type I or type II. We compare these results to the type I genetic architecture known from the independently evolving species pair, *L. kohalensis* and *L. paranigra,* which diverged in pulse rate within a single island, that is, Hawaii [31,32,54]. Although *L. cerasina* (Hawaii Island) likely arose as a consequence of an interisland speciation event from the ancestral source range (Maui) of *L. eukolea* [53], we hypothesize a type I genetic architecture, based on previously published biometrical results [39].

The second aim of this study, inspired by theoretical predictions [22] as well as recent findings on the genetics of mating behavior [25–27], is to test the hypothesis that the functionally related genes cluster in QTL regions, thereby structuring the potential for adaptive change. Using genome-wide, functional genetic data from both sexes, and across ontogenetic stages and reproductive states, we annotate the *Laupala kohalensis* genome [55] and perform gene set enrichment analysis. Based on neurobiological insights into cricket song generation and neurogenetic insight into song variation in *Drosophila*, we expect QTL regions to be enriched for genes with putative functions in neuromuscular processes associated with song production, rhythmic behaviors, or mating behavior. This would suggest that “pools” of functionally related genes associate with QTL regions and thus provide a large source of potential genetic variation for phenotypic change (as opposed to QTL regions harboring single genes that would limit potential evolutionary contributions to the observed phenotypic effect).

## 2. Materials and Methods

*Laupala eukolea* nymphs were collected in 2012 in Kipahulu Valley on Maui at Ginger Camp (20° 41’ 60.000’’ N; 156° 5’ 18.000’’ W) and Palikea Peak (20° 40’ 20.640’’ N; 156° 4’ 5.160’’ W); *L.cerasina* were c°llected in Kalopa State Park (20° 2’ 13.200’’ N; 155° 26’ 36.960’’ W) on Hawai’i Island. Animals were kept in the lab in plastic cups under constant temperature (20°C) and humidity, and were provided cricket chow (Fluker Farms, LA) *ad libitum* as well as substrate to lay eggs. Males and females were kept separately to ensure virginity of all animals. They were phenotyped between three and ten weeks after the final molt, during which sexual receptivity is maximized in *Laupala* [38]. Two families of first generation interspecific hybrids (HC1 & HC2) were each generated by mating a *L. cerasina* male with a *L. eukolea* female. Several males and females from each family (14 full sib pairs from each of the families) were used to obtain second generation hybrids.

Male songs were recorded following the methods in Shaw (1996) [35]. A single recording for each individual (*L. cerasina*, n = 24; *L. eukolea*, n = 16; F2, n = 230) was made between 10 A.M. and 4 P.M. Virgin, adult males were recorded individually in a plastic container with screen covers in an anechoic and temperature-controlled chamber using a SONY Pro Walkman cassette recorder and SONY microphone. Songs were then digitized using SOUNDSCOPE / 16 (GWI Instruments, Cambridge, MA, USA) at 44.1 kHz to generate an oscillogram displaying trains of pulses (singing bouts). We estimated the pulse rate by averaging the inverse of five pulse periods (measured from the onset of a pulse to the onset on the next pulse) measured from a single singing bout.

We extracted DNA from whole adult male crickets using the DNeasy Blood & Tissue Kits (Qiagen, Valencia, CA, USA). Genotype-by-Sequencing library preparation and sequencing were done in 2014 at the Genomic Diversity Facility at Cornell University following [56]. The *Pst* I restriction enzyme was used for sequence digestion and DNA was sequenced on the Illumina HiSeq 2000 platform (Illumina Inc., USA) with 100 bp single end reads.

Reads were trimmed and demultiplexed using Flexbar v2.5 [57] and then mapped to the *L. kohalensis de novo* draft genome using Bowtie2 v2.2.6 [58] with default parameters. We then called SNPs using two different pipelines: The Genome Analysis Toolkit v3.6.0 (GATK) [59,60] and FreeBayes v0.9.13 [61]. For GATK we used individual BAM files to generate gVCF files using ‘HaplotypeCaller’ followed by the joint genotyping step ‘GenotypeGVCF’. We then evaluated variation in SNP quality across all genotypes using custom scripts in R v3.3.1 [62] to determine appropriate settings for hard filtering based on the following metrics (based on the recommendations for hard filtering section “Understanding and adapting the generic hard-filtering recommendations” at https://software.broadinstitute.org/gatk/ accessed on 28 February 2017): quality-by-depth, Phred-scaled *P*-value using Fisher’s Exact Test to detect strand bias, root mean square of the mapping quality of the reads, u-based z-approximation from the Mann-Whitney Rank Sum Test for mapping qualities, u-based z-approximation from the Mann-Whitney Rank Sum Test for the distance from the end of the read for reads with the alternate allele. For FreeBayes we called variants from a merged BAM file using standard filters. After variant calling we filtered the SNPs using ‘vcffilter’, a Perl library part of the VCFtools package [63] based on the following metrics: quality (> 30), depth of coverage (> 10), and strand bias for the alternative and reference alleles (SAP and SRP, both > 0.0001). Finally, the variant files from the GATK pipeline and the FreeBayes pipeline were filtered to only contain biallelic SNPs with less than 10% missing genotypes using VCFtools v0.1.15. We retained all SNPs that had identical genotype calls between the two variant discovery pipelines. We then pruned our data further to contain only ancestry informative markers (i.e. grandparents are homozygous for alternative alleles), one SNP per scaffold, markers with no or only limited segregation distortion from the expected 1:2:1 (autosomal) and 1:1 (X-linked) ratios (χ^2^-square associated q-value ≤ 0.05, i.e. a 5% false discovery rate, [64]), and markers with fewer than 99% of their genotypes in common with other marker loci (i.e. exclude one of each pair of marker loci with identical genotypes for all individuals).

For each of the hybrid cross families, HC1 and HC2, we created linkage maps separately using MapMaker v3.0b [65] taking the following three steps. First, markers were grouped into linkage groups using the ‘group’ command with the ‘default linkage criteria’ set to 4.0 (LOD, logarithm of the odds) and 35 (cM). Second, for each group, a subset of the marker passing a series of quality criteria, i.e. informative, well-spaced markers, were ordered using the ‘order’ command. Informative markers were those with no missing genotypes and more than 2.0 cM apart from other markers. Marker order was compared with regression mapping in JoinMap v4.0 [66] and inconsistencies were resolved by minimizing stress (in JoinMap) and map length and maximizing the likelihood (in MapMaker). Third, remaining markers were added with the ‘build’ command and order was verified using the ‘ripple’ command. At this step, markers were added to the map satisfying a log-likelihood threshold of 4.0 for positioning of the marker (i.e., the assigned marker position is 10,000 times more likely than all other positions), then adding remaining markers at a log-likelihood threshold of 3.0, followed by a final addition at a log-likelihood threshold of 2.0. Subsequently, any unincorporated markers were discarded. To determine the final marker order we used the ‘ripple’ command with a window size of 6 markers and a log-likelihood threshold of 2.0. Arbitrary orders in marker dense regions (i.e., orders with similar likelihoods) were resolved using information from both HC1 and HC2 maps, choosing the order that maximized the likelihood and minimized the map length (measured in cM) for both cross families.

Finally, we merged the separate HC1 and HC2 maps using the R package LPmerge v1.6 [67]. LPmerge uses linear programming to combine two maps from independent populations based on the similarities in marker order. Incongruent marker orders between HC1 and HC2, i.e. linear inequalities, were solved based on the weight assigned to each independent linkage map. The solution also depended on the size of the interval, *K*, in which conflicting markers were detected and re-ordered (or removed if no solution was found and removing a constraining marker improved the linear equality). For each linkage group separately, we varied the weighting of the two linkage maps and the interval in which the linear inequality was resolved (*K*) to find the consensus map associated with the lowest mean and variance of the root-mean-squared error between the consensus map and the original maps.

We used composite interval mapping (CIM) and multiple-QTL models (MQM) in the R/qtl v1.42 [68] package to detect and locate QTL and calculate effect sizes independently for HC1, HC2, and for the merged (consensus) map. We first performed a single QTL scan using the scanone function with the multiple imputation method [69] and the Haley-Knot” regression [70]. For CIM, we then ran a model using the Haley-Knot regression in 20 cM windows, with the number of included marker covariates dependent on the number of QTL detected in the single QTL scan. We then performed a two-dimensional QTL scan using the Haley-Knott regression to detect pairs of QTL and interaction effects among QTL and permuted the two-dimensional QTL to establish penalized likelihood criteria for main and interaction effects. We subsequently built a multiple QTL model, starting with the QTL with the highest LOD score in the single QTL scan, refining the position using ‘refineqtl’, and then scanning for additional QTL. We continued adding QTL (followed by refining their position) until an additional QTL did not improve the LOD score of the model beyond the penalized LOD score threshold for main effects at the alpha = 0.05 level. After the addition of each QTL, we checked for potential QTL interactions that would improve the multiple QTL model beyond the (heavy) penalized LOD score threshold for interaction effects at the alpha =0.05 level. Finally, we estimated effect sizes by fitting the QTL model and using the drop-one-term analysis. In the merged map, we included cross type as a covariate in all steps described above.

To estimate the true number of genetic loci underlying pulse rate variation based on the QTL results, we use the method of Otto and Jones [71]. We estimated the minimum detectable QTL effect size using equation 11 in [71], specifying *a*_*min*_ as the smallest QTL we detected in the QTL scan (for HC1, HC2, or for the combined map). We then used equation 6 in [71] to estimate the true number of loci. The calculations were done using custom functions in R.

To look for candidate genes in QTL regions and test for enrichment of specific gene functions we first assembled the transcriptome to obtain information about putative gene function of the loci within regions of interest. First generation, lab-reared *L. cerasina* individuals were used for RNA-sequencing and transcriptome assembly. A total of ten samples were stored in RNAlater following the manufacturers recommendations (Qiagen, Germany) and pooled by sex prior to sequencing: four adult males (two sampled in the morning and two sampled in the evening, one of which was sampled immediately *after* mating), four adult females (likewise, two were sampled in the morning and two sampled in the evening, one of which was sampled immediately *after* mating), and a juvenile male and female. The tissue was homogenized using sterilized forceps in RNAlater. RNA was extracted using the RNAeasy kit (Qiagen, Germany). A quality check was done using a NanoDrop spectrophotometer (Thermoscientific, Wilmington, DE, USA) and the Agilent Bioanalyzer 2100 [72]. The samples were then sequenced on a single lane on the Illumina HiSeq 2000 platform with 50 bp paired-end reads. Reads were processed using Fastq-mcf from the Ea-Utils package [73] with the parameters -q 30 (nucleotides from the extremes of the read with qscore below 30 were trimmed) and -l 30 (reads with lengths below 30 bp discarded). Read duplications were removed using PrinSeq v0.20.4 [74] and reads were corrected using Musket v1.0.8 [75] with the default parameters.

We assembled the *L. cerasina* transcriptome using Trinity’s v.2.4.0 [76] genome-guided assembly pipeline. We used the *Laupala kohalensis* reference genome [55] and a maximum intron size cut-off of 5,000 bp. We first created a HISAT v4.8.2 [77] index of the genome and then aligned the male and female paired-end reads to the genome using default settings. We sorted the resulting alignment SAM file and converted it to BAM format using Samtools v1.5 [78]. We then did the genome-guided assembly using Trinity and checked the quality of the assembly by calculating N50 statistics, mapping the male and female reads back to the transcriptome using Bowtie2, and searching for conserved eukaryote and arthropod genes from the BUSCO database v.2.0.1 [79]. We then mapped the reads back to the transcriptome using GMAP v 2017-05-08 [80] and BLAT v35×1 [81], and used custom R scripts to retain a single best hit scaffold for each transcript based on the coverage, identity, and number of matched bases.

We checked for and removed contaminants using NCBI’s VecScreen, using the UniVec Core database and the recommended blastn parameter values https://www.ncbi.nlm.nih.gov/tools/vecscreen/about/, accessed July 20 2017). We then functionally annotated the transcripts in three steps using BLAST [82,83]: first we matched our transcripts against the *Drosophila melanogaster* proteome using an e-value cut-off of 1e-5. Any transcripts that were not assigned a putative *D. melanogaster* homolog were matched (at the same threshold) with the Uniprot/Swissprot data base [84], limiting our search to arthropod proteins. Finally, we matched any remaining transcripts after second step against the non-redundant protein database at NCBI, with an e-value cut-off of 1e-5 and limiting our search to animal proteins.

We then used hmmer2go v3.1 (https://github.com/sestaton/HMMER2GO) to estimate Open Reading Frames (ORFs) and translated only a single, longest ORF per transcript. We annotated the retained protein sequences using InterProScan v5 [85,86]. We imported the transcriptome FASTA file, the XML output from the BLAST searches, and the InterPro results into Blast2Go v4.1.9 [87]. We recovered the original BLAST best hit and ran the Gene Ontology mapping using default settings. We then merged all these results and ran the Annotation tool.

If genes controlling pulse rate variation in *Laupala* are clustered in specific genomic regions rather than distributed randomly across the genome, we expect QTL regions to contain several (putative) causal genes for interspecific pulse rate variation. We first used topGO v2.24.0 from R’s BioConductor v3.3 environment [88] for gene set enrichment analysis. We combined all the transcripts matching scaffolds within the peak +/− 1 LOD interval for each of the seven QTL peaks in the consensus linkage map (i.e. scaffolds with a marker linked to pulse rate variation at a likelihood of no less than the peak LOD score minus 1). We then used the parent-child p-value correction [89] and an additional false discovery correction [64] with the ‘p.adjust’ function in R (GO terms were considered enriched at a false discovery rate of 10% or less). As these analyses are potentially confounded by pseudo-replication in the transcriptome assembly (e.g., due to varying occurrences of exonic splice sites and transcription start sites), we perform the above analysis after collapsing all the transcripts that mapped to the same scaffold and were considered putative alternative splice form based on their annotations (i.e. transcripts that have the same predicted gene product, are annotated with different isoforms of the same gene or partial and full hits of the same gene). For comparison, we also conduct the GO enrichment analysis using the original annotated transcriptome. Finally, we manually inspected putative gene function in QTL regions by examining experimentally proven biological and molecular functions for the highest ranked BLAST annotations on FlyBase [90].

## 3. Results

### 3.1 Linkage mapping

We mapped a total of 298 and 416 markers for HC1 (n=94) and HC2 (n=136), respectively to 8 linkage groups corresponding to the seven autosomes and the X chromosome in *Laupala* (Fig S1). The map lengths were 821.2 cM and 734.1 cM corresponding to 2.72 cM and 1.76 cM, per marker, respectively. Merging the maps resulted in 508 unique markers at a total map length of 776 cM (1.52 cM/marker). The maps are broadly similar (in order and length) across the two mapping families. One linkage group, LG1 shows substantially higher recombination rates among the markers in HC1 relative to HC2 (Fig S1), which may be due to sampling variance, structural variation, or both. Comparing the marker order on LG1 with the homologous LG in two interspecies crosses [55] revealed that this region is likely inverted in the map for HC1 (Fig S2), indicating that a large pericentric inversion is segregating in either *L. cerasina, L. eukolea*, or both.

### 3.2 QTL mapping

*Laupala cerasina* and *L. eukolea* males have non-overlapping, normally distributed pulse rate distributions (Fig 1). The mean pulse rate difference was 1.66 pulses per second (pps; Table 1). The F2 song phenotype was normally distributed and the means of the HC1 and HC2 progeny did not differ significantly (*P* = 0.0724). Considering the joint F2 distribution, the slow tail of the distribution partially overlapped with the *L. cerasina* phenotypic distributions although the fast tail did not overlap the *L. eukolea* phenotypic distribution (Table 1, Fig 1).

**Table 1.** Phenotypic distributions. The mean and standard deviation of the pulse rate (pulses per second) and the sample size are shown for the parental species and the F2 generation (both cross types, HC1 and HC2).

|  | Mean<br>(pps) | sd | n |
| --- | --- | --- | --- |
| *L. cerasina* | 2.33 | 0.07 | 24 |
| *L. eukolea* | 3.99 | 0.12 | 16 |
| Environmental<br>Variance* | 0.02 | 0.13 |  |
| F2 HC1 | 3.11 | 0.20 | 94 |
| F2 HC2 | 3.16 | 0.23 | 136 |
| F2 mean | 3.14 | 0.22 | 230 |

\*Environmental variance was calculated following [91]; “mean” in this context refers to the environmental variance, while “sd” is the square root of that variance.

**Figure 1.**
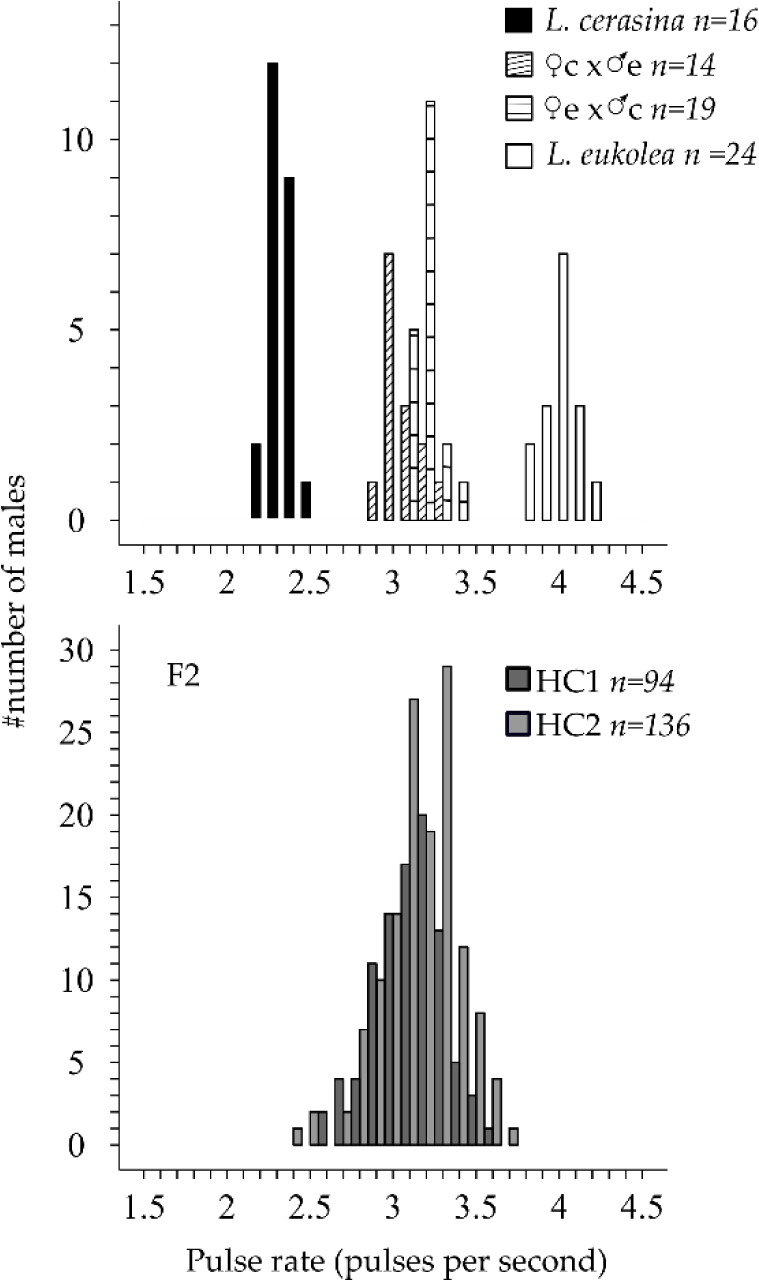
Phenotypic distributions for the parental lines and first and second-generation hybrid offspring. The data from the F1 hybrids in the top panel are from Oh et al. 2012 [39] and are shown to illustrate F1 distributions as well as the maternal bias in pulse rate inheritance. The F2 data are split by cross type (HC1 and HC2).

Using CIM and MQM, we detected two moderate effect (˜10% of the species difference) QTL on LG1 and LG3 (Fig 2, Table 2) in both the HC1 and HC2 mapping populations. We note that the LOD profile for MQM in HC1 suggests an additional peak on LG1 (at 37 cM), but neither CIM, nor MQM supports this. We believe the “phantom” peak derives from the putative inversion creating genotype-phenotype associations for markers outside the QTL region in some individuals but not in others. In HC2, we additionally detected two smaller effect QTL (< 5%), on LG5 and LGX. Merging the maps and combining the sample sizes also revealed small effect QTL on LG2, LG4, and LG7 (Table 3), using MQM (Fig 2). When adding these additional QTL to the MQM model, the peak on LG3 shifts approximately 10cM posteriorly, but the 1-LOD intervals of the former and refined QTL overlap (Fig S1), and consequently does not impact our annotations (see below).

**Table 2.** QTL results from HC1 and HC2. QTL were mapped using the maps for the 94 HC1 and 136 HC2 F2 individuals. LOD: log-of-odds. A and B alleles denote *L. cerasina* and *L. eukolea* alleles, respectively. All QTL effects (in pulses per second) are significantly different from zero. ESD: environmental standard deviation.

| effects (in pulses per second) are significantly different from zero. ESD: environmental standard deviation. |  |  |  |  |  |  |  |  |  |  |  |
| --- | --- | --- | --- | --- | --- | --- | --- | --- | --- | --- | --- |
| Linkage group | Location (cM) | LOD | Nearest scaffold | Marker location | Phenotypic Value |  |  | Effect (pps) | % parental difference | #ESDs | % F2 variance |
|  |  |  |  |  | AA | AB | BB |  |  |  |  |
| HC1 |  |  |  |  |  |  |  |  |  |  |  |
| 1 | 119 | 17.81 | S004794 | 119.4 | 2.88 | 3.14 | 3.31 | 0.220 | 13.23 | 1.75 | 50.60 |
| 3 | 72.0 | 7.04 | S001552 | 71.3 | 2.95 | 3.13 | 3.16 | 0.116 | 6.95 | 0.92 | 14.96 |
| HC2 |  |  |  |  |  |  |  |  |  |  |  |
| 1 | 56.9 | 23.27 | S002490 | 56.9 | 2.90 | 3.18 | 3.31 | 0.217 | 13.04 | 1.72 | 38.67 |
| 3 | 65.0 | 13.92 | S000355 | 67.1 | 3.03 | 3.17 | 3.28 | 0.148 | 8.90 | 1.08 | 19.42 |
| 5 | 38.4 | 5.55 | S002808 | 38.4 | 3.04 | 3.17 | 3.25 | 0.074 | 4.45 | 0.60 | 6.67 |
| X | 32.0 | 3.42 | S000108 | 29.7 | 3.13 | -<sup>a</sup> | 3.19 | 0.049 | 2.95 | 0.39 | 3.96 |

<sup>a</sup> cricket males are hemizygous

**Table 3.** QTL results from the combined map. QTL were mapped using the consensus map for the 230 F2 individuals. LOD: log-of-odds. A and B alleles denote *L. cerasina* and *L. eukolea* alleles, respectively. All QTL effects are significantly different from zero. ESD: environmental standard deviation.

| Linkage group | Location (cM) | LOD | Nearest scaffold | Marker location | Phenotypic value |  |  | Effect (pps) | % parental difference | #ESDs | % F2 variance |
| --- | --- | --- | --- | --- | --- | --- | --- | --- | --- | --- | --- |
|  |  |  |  |  | AA | AB | BB |  |  |  |  |
| 1 | 59.0 | 40.73 | S001131 | 58.8 | 2.89 | 3.17 | 3.31 | 0.203 | 12.22 | 1.61 | 36.39 |
| 2 | 71.0 | 4.78 | S001921 | 69.1 | 3.05 | 3.15 | 3.18 | 0.055 | 3.30 | 0.44 | 2.90 |
| 3 | 79.9 | 21.28 | S000385 | 79.9 | 3.02 | 3.14 | 3.23 | 0.123 | 7.40 | 0.98 | 15.34 |
| 4 | 59.8 | 3.97 | S016452 | 59.0 | 3.10 | 3.12 | 3.23 | 0.053 | 3.19 | 0.42 | 2.39 |
| 5 | 36.9 | 8.32 | S002445 | 36.9 | 3.05 | 3.16 | 3.19 | 0.068 | 4.08 | 0.54 | 5.23 |
| 7 | 5.0 | 3.51 | S007011 | 9.8 | 3.08 | 3.14 | 3.20 | 0.045 | 2.68 | 0.35 | 2.10 |
| X | 38.0 | 5.46 | S003132 | 37.2 | 3.10 | - | 3.17 | 0.041 | 2.46 | 0.32 | 3.34 |
| cross | - | 2.24 | - | - | - | - | - | 0.092 | 5.53 | 0.44 | 1.33 |

**Figure 2.**
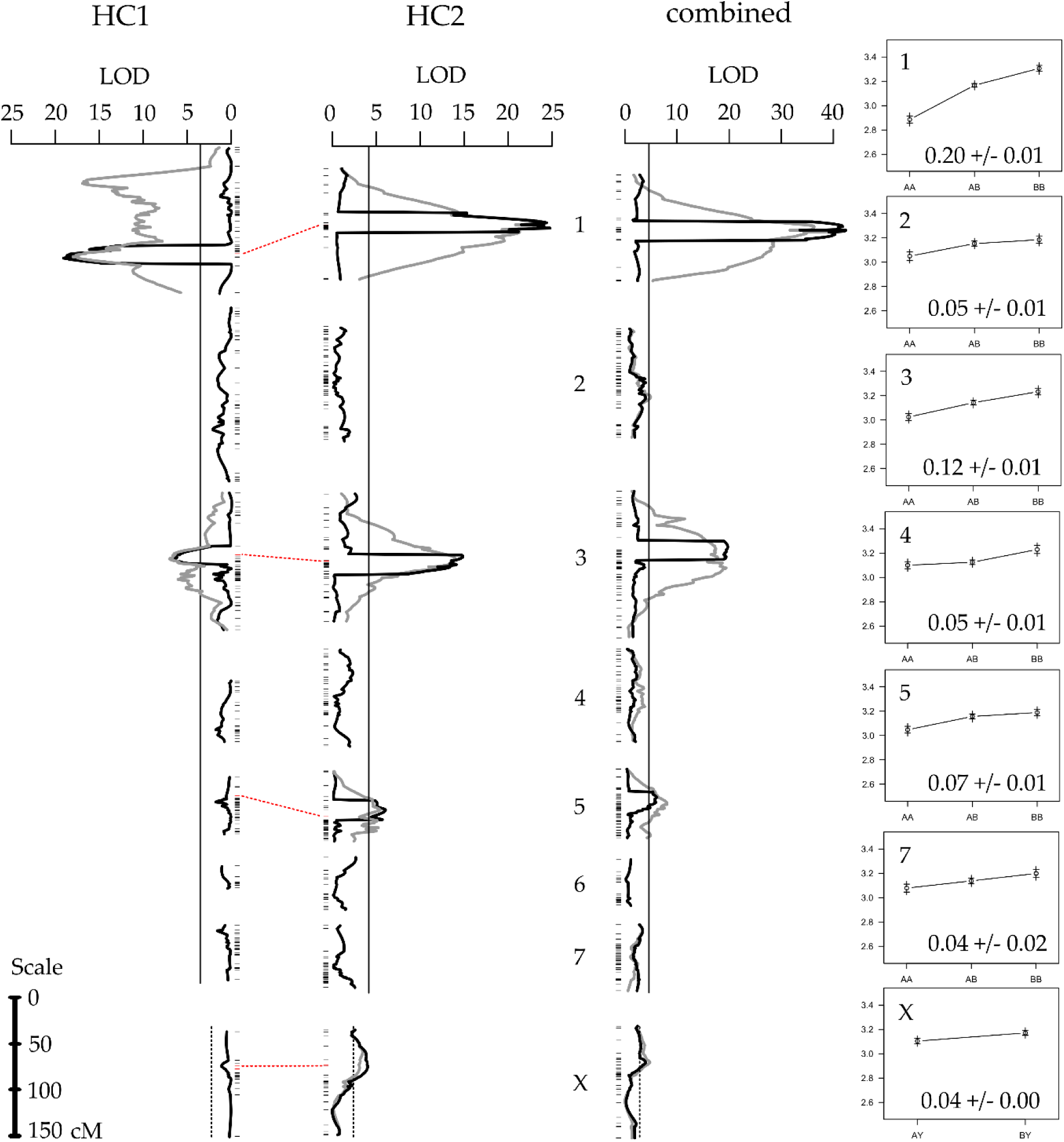
QTL scan. Results from composite interval mapping (black lines) and multiple QTL models (grey lines, only for linkage groups with significant QTL) The vertical solid and dotted lines show the experiment-wide 5% significance threshold for CIM for autosomes and the X-chromosome, respectively. Between HC1 and HC2, horizontal dotted lines connect homologous markers associated with QTL peaks (within the CIM windows) to indicate the overlap between the QTL scans in the different mapping families (see Fig S1 for more detail). The panels on the far right show the effect size of each of the QTL as the pulse rate mean +/− standard error for each of the genotype categories AA (left), AB (center), BB (right).

All (haploid) QTL effects were significantly larger than zero (P < 0.05) and of the same sign (Table 2, Table 3). Together, the seven QTL for the combined HC1 and HC2 mapping populations explained 35.41% (or 0.59 pps) of the haploid phenotypic difference between the parental lines (Table 3; 20.22% and 31.71% for HC1 and HC2, respectively, Table 2). None of the QTL had significant dominant effects (Table S1) and no interactions between additive QTL were detected, thus the total amount (twice the additive haploid effect) of the pulse rate difference explained by the seven QTL is 70.82% (or 1.16 pps) of the species difference. The QTL on LG1 and LG3 mapped to approximately the same location in HC1 and HC2, with the marker nearest to the peak in HC2 directly flanking the peak marker in HC1 (Fig 2, Table S2).

Based on the MQM results for HC1 and HC2 and using the method in [71] we estimated the true number of loci to be 9.55 (95% confidence interval = [1.59-29.49]; detection threshold, θ = 0.08 pps) and 9.4 (95% CI = [2.93-21.86]; θ = 0.03 pps), respectively. Using all 230 F2 individuals, we estimated the true number of loci at 16.6 ((95% CI = [7.14-32.12]; θ = 0.04 pps). The variation in the true number of loci between HC1, HC2, and the combined sample reflects variation in the mean across all detected QTL effects as well as in the lowest detected QTL effect, which is dependent on the sample size.

### 3.3 Transcriptome assembly and annotation

We used the genome-guided assembly from the Trinity pipeline [76] to assemble the transcriptome. Of the 50,148,157 reads after filtering (52,980,661 prior to filtering) that were used to assemble the transcriptome, 90.58% mapped to the *Laupala kohalensis* reference genome. The assembly had a total length of 53,928,392 bp, the median contig length was 397 bp, and the contig N50 was 1,805 bp. Mean coverage was 49.9x and 40.8x for female and male reads, respectively. The male and female reads mapped back to the transcriptome with high confidence, at mapping rates of 92.05% and 92.26%, respectively. The BUSCO analysis indicated that we captured a large amount of conserved eukaryote and arthropod genes with 97.4% and 95.2% complete BUSCO hits, respectively. Using the *D. melanogaster* proteome, the arthropod specific Uniprot/Swissprot database and NCBI’s non-redundant database of animal proteins, we successfully annotated 19,713 transcripts (32.2% of all transcripts) at a combined length of 32,039,513 bp (59% of the full assembly). A total of 17,577 transcripts have a Gene Ontology (GO) annotation.

### 3.4 Gene set enrichment analysis

There were 179 scaffolds within 1 LOD of the seven QTL peaks combined. After collapsing putative alternative splice forms of genes (transcripts on the same scaffold with identical predicted protein products or annotated with different isoforms of the same gene), we mapped a total of 1,298 annotated transcripts to 171 of the 179 scaffolds (Table S3). We tested whether QTL regions were significantly enriched for biological processes that are relevant to cricket mating behavior, i.e. sexual (acoustic) communication, muscle contraction and pacemaker genes, and various neuromuscular properties and neuromodulators of rhythmic behaviors. We find significant (FDR of corrected p-value < 10%) enrichment of 37 biological processes, many of which are related to neurobiological and muscular development, i.e. peripheral nervous system development, dendrite guidance, brain morphogenesis, and neuromuscular junction development in the combined set of all 7 QTL regions (Fig 3, Table S4). Similarly, for each of the seven QTL regions separately we find significant enrichment of, among others, central complex and motor neuron development (LG1), hormonal and pheromonal biosynthetic pathways (LG2), neurotransmitter transport and mating behavior (LG3), peripheral nervous system development (LG4), flight and locomotor behavior (LG5), neuromuscular junction development (LG7), and calcium ion transport (LG X; Table S5). Using the 1-LOD interval for the QTL on LG3 prior to the 10 cM shift in the QLT peak gave identical results for the enrichment of that QTL region. Excluding the QTL on LG7, which has a large confidence interval (and only weak phenotypic effect), also gave similar results; although, locomotory behavior is no longer significantly enriched (data not shown).

**Figure 3.**
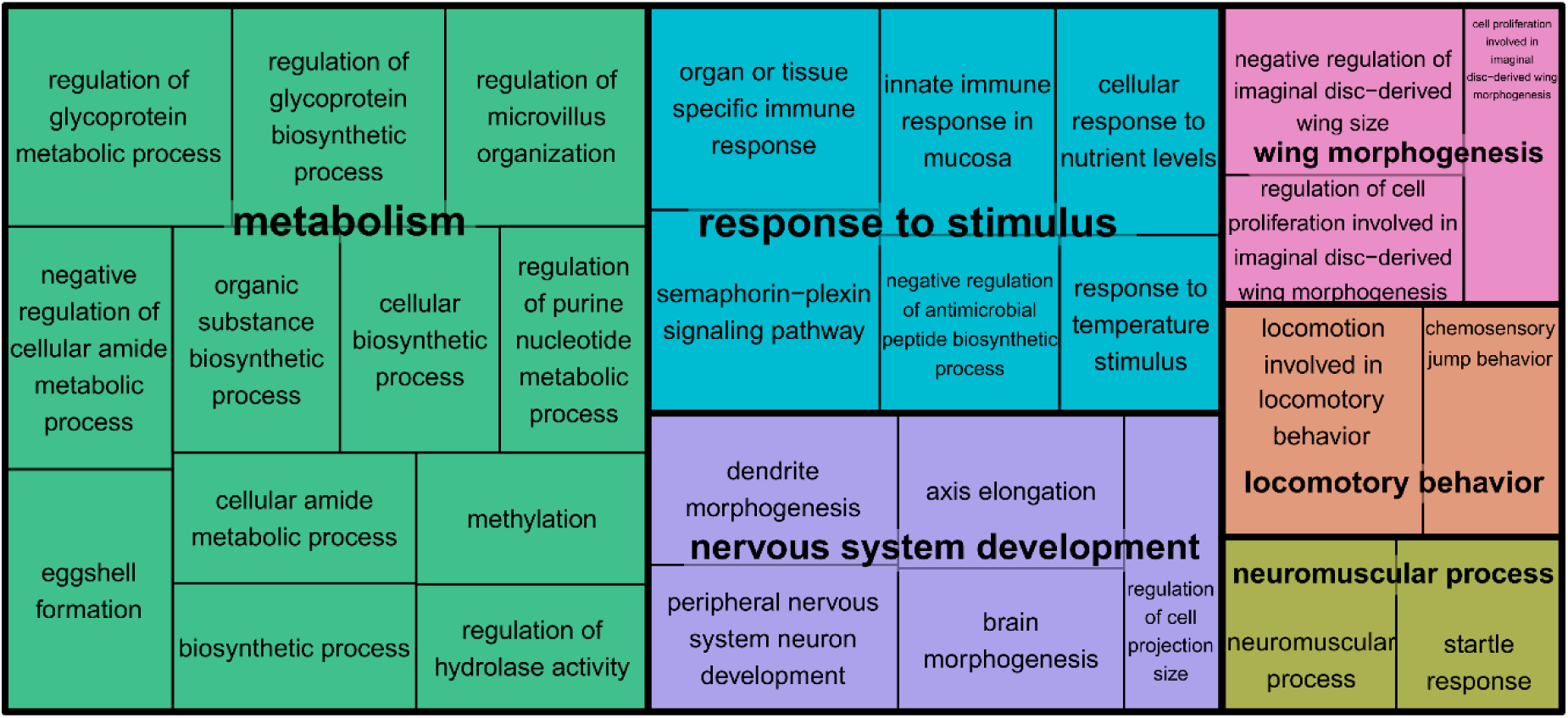
Treemap of enriched GO categories. The GO terms were subset removing all redundant GO terms in REVIGO [92] at the medium-similarity criterion (0.7) are grouped based on the taxonomic relations among GO terms. Colors connect GO terms belonging to the same cluster. The size of the panels scales with the negative 10-logarithm of the p-value for the enrichment test.

When putative alternative splice forms are not collapsed into a single annotated gene product (i.e. the original annotation of the transcriptome assembly), 1,724 annotated transcripts map to scaffolds within the 7 QTL intervals (Table S6). We find substantially more enriched GO terms (227 biological processes). However, many of the same terms are enriched compared to the analysis above, both for the overall enrichment (Table S7), as well as for the linkage groups separately (Table S8): e.g. terms related to neuromuscular development (LG1, LG4), development, maintenance, and transmission at synapses (LG1, LG3, LG5, LGX), rhythmic and locomotor behaviors (LG4, LG7), mating behavior (LG3), hormone and pheromone production (LG2), and nervous system development (LG4, LGX; Table S8).

## 4. Discussion

Behavioral isolation is often an important barrier to gene flow in the earliest stages of animal speciation, but we know little about the number and distribution of underlying genetic loci and their relationships with the tempo and mode of evolution. In island systems where founder effects have been hypothesized in the history of species radiations, shifts in the phenotypic and genetic environments may catalyze speciation through the interactions of genetic drift and the genetic architecture of traits [11,93], in particular for traits involved in sexual isolation [93–95]. With a type II genetic architecture (epistatic, major gene/modifier), the genetic system becomes reorganized during the genetic upheaval following the founder event, and thus acts as a lynch pin for speciation.

Here, we show that the genetic architecture of a major premating barrier in *Laupala* speciation following island colonization has a type I (polygenic, additive) genetic architecture, as is common for sexual signals [3], rather than a type II genetic architecture that can promote founder effect speciation. This finding, compared to previous intra-island divergence events, indicates that similar genetic architectures underlie repeated episodes of mating song divergence in *Laupala*, independent of biogeographic history. A tentative explanation for the observed similarity may be that QTL regions associate with functionally related genetic clusters, within which a large pool of genes and numerous functional sites could potentially contribute to song evolution, thus restricting the spatial genomic regions of phenotypic change but not the number of quantitative trait nucleotides. Within QTL regions, we also identify many potential homologs of genes implicated in *Drosophila melanogaster* courtship behavior and in various neurophysiological processes that might be important for song rhythm divergence in *Laupala*.

### 4.1 The genetic architecture of interspecific pulse rate divergence

Our data support the hypothesis that song divergence between *L. eukolea* and *L. cerasina* is associated with a Type I polygenic genetic architecture. We detected seven small-to-moderate effect QTL for pulse rate divergence, six QTL each on different autosomes and an additional small effect X-linked QTL. There were no detectable interaction effects among these identified QTL. The position and effect size of the two largest QTL between replicate families were largely the same (sharing the position of peak or flanking (or both) markers on LG1 and LG3; Table S2). The effects of sample size being well-known [96], we merged the families to leverage the power of increased sample size. We found the position of the two QTL detected in HC2 but not in HC1 (on LG5 and LGX) shared common positions and effect sizes with this combined analysis (Table S2). In the combined analysis, we further detected three small effect QTL on LG2, LG4 and LG7. Adding all QTL terms to the MQM model resulted in a refined estimate of the peak QTL location on LG3 in the combined map. Further exploration revealed that this position refinement was not specific to the map used in the analysis, as simulated QTL experiments as well as adding additional QTL (that are borderline significant or only “suggestive”) to the HC1 and HC2 models yield similar effects (not shown). Drawing on all QTL identified, and the method of Otto and Jones [71], we estimate that the true number of loci is likely more than ten.

### 4.2 Inter versus intra-island speciation

Contrasting the results for *L. cerasina* and *L. eukolea* with those for the intra-island species pair *L. kohalensis* and *L. paranigra* [54] suggests that the genetic differentiation of song has occurred by similar genetic architectures in these independent divergence events: we detect similar numbers of loci on the same linkage groups with comparable effect size distributions (Table 4). Morphological [32] and molecular evidence [31] place the replicate species pairs considered above in independent species groups. Based on the young age of the Big Island [97], to which *L. kohalensis, L. paranigra* and *L. cerasina* are endemic, both these species pairs have likely diverged in the last 500,000 years [31], but differ in the biogeographic context of speciation. We estimate that the two major QTL, on LG1 and LG3, explain around 12% and 8% of the average phenotypic difference between *L. eukolea* and *L. cerasina*. The homologous linkage groups in the genetic map of *L. kohalensis* and *L. paranigra* (linkage group numbers are the same) have QTL explaining around 9% and 10% of the parental difference, respectively. Likewise, both the present study and the *L. kohalensis* x *L. paranigra* study find additional QTL on LG4, LG5 and the X chromosome, with no detected interactions among loci. Previously, biometric studies had also revealed multiple independent genetic factors and an X-effect [35,39]. Moreover, phenotype associations on LG6 and LG7 were weak or absent in both studies and, further, estimates of the true number of loci are > 10 in both studies, suggesting a polygenic architecture for both inter-island and intra-island speciation. Finally, all QTL effects estimated in this study and in Shaw et al. (2007) are of the same sign, consistent with a hypothesis of directional selection [98]; that is, alleles from the fast species increase the pulse rate, whereas alleles from the slow species decrease the pulse rate of F2 hybrids. Thus, overall these results support similar genetic architectures for pulse rate divergence regardless of biogeographic context. We caution, however, that other phenotypes may or may not follow this pattern, e.g. cuticular hydrocarbon variation [99], and merit further investigation.

**Table 4.**
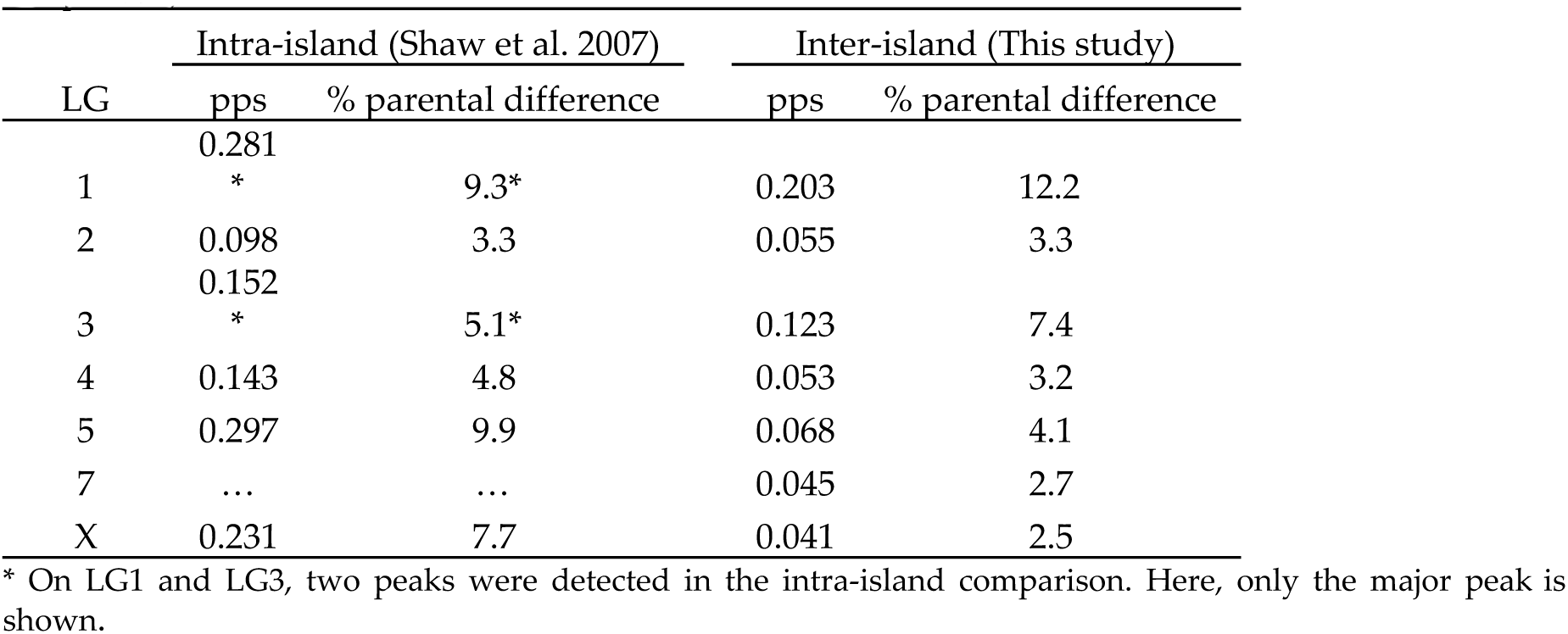
Comparison of QTL effect for intra-island versus inter-island divergence. The QTL effects (in pulses per second and as a percentage of the parental difference) are shown for the Shaw et al study (the intra-island comparison, results from Table 2b in that study) and for the results of the present study (the inter-island <u>comparison</u>).

It should be acknowledged that there are also some differences between the two species pairs. The pulse rate difference between *L. cerasina* and *L. eukolea* is roughly half of that between *L. paranigra* and *L. kohalensis* [54]. We detect just seven QTL in the present study as opposed to eight QTL in the *L. paranigra* and *L. kohalensis* cross, despite similar sample sizes and much denser genotype sampling in the present work. We also find smaller average effect sizes for *L. cerasina* and *L. eukolea* QTL. For example, while the QTL effects on LG4 and LG5 are present in both species pairs, they are smaller (both in absolute and relative terms) in the *L. cerasina* and *L. eukolea* cross than the *L. kohalensis* x *L. paranigra* cross (0.05 and 0.07 pps versus ˜0.14 and ˜0.29 pps, respectively). Additionally, we did not find evidence for minor QTL on LG1 and LG3 in the present study although both linkage groups were found to harbor minor peaks in the 2007 study. It must be noted that the resolution at which we resolve QTL regions is in the order of several centi-Morgans. Additionally, Shaw et al. (2007) used AFLP markers while we use GBS markers. Therefore, comparisons of the genetic architectures between these two independent species pairs can only be made at the chromosomal (linkage groups) level because we lack sequenced-based markers and precise genomic locations of QTL in *L. paranigra* x *L. kohalensis*. More detailed information is needed to test whether independent divergence in pulse rates is associated with convergent genetic mechanisms.

However, the overall similarity in genetic architecture is significant in that *Laupala* has one of the fastest rates of speciation known, yet a type I genetic architecture underlies the interspecific genetics of an important speciation phenotype in this radiation, independent of the biogeographic context. Moreover, our findings suggest that differences in the extent of phenotypic differentiation are due to larger effect sizes of substitutions in the same QTL regions rather than that additional QTL are involved in the more diverged species pair. Further comparative work is needed to probe the generality of these findings and to better illuminate the relationship between biogeography, magnitude of phenotypic divergence, and genetic architecture of speciation. Additionally, potential mechanisms that constrain the number of possible locations in the genome where genetic changes that contribute to song rhythm variation can occur need to be examined. One potential mechanism could be that causal loci are not randomly distributed across the genome, but instead cluster in specific genomic regions.

### 4.3 Behavioral gene clusters

Clustering of causal loci that are important to reproductive isolation is expected on both theoretical [22,100] and empirical [25–28] grounds and can have dramatic consequences for the mode and rate of evolution. If the genetics of behavioral isolation in *Laupala* is characterized by the clustering of causal loci, we expected to find strong enrichment of genes with putative roles in cricket song and mating behavior in QTL regions. In line with these expectations, we find enrichment of GOs that can be tied to the (neuro)biology of cricket mating behavior. Enrichment is evident for all QTL combined, with or without the QTL on LG7 (which has an exceptionally broad confidence interval). Moreover, the pattern is not driven by a single region, but rather, significant enrichment contributions derive from every QTL region separately. Interestingly, some of these QTL fall in regions of low recombination (e.g. QTL on LG1, LG3, and LG5), in the central parts of the chromosomes [55] where we observed high marker densities (Fig S1). Reduced (interspecific) recombination rates can reinforce linkage disequilibrium between co-adapted loci over larger genomic distances. Together, these findings suggest that acoustic mating behavior divergence in crickets is potentially associated with clusters of causal loci rather than randomly distributed loci.

Overall, the finding that QTL regions are strongly enriched for homologs of genes involved in neuromodulation and nervous system development is an exciting novelty in the attempt to unravel the genetic architecture of premating isolation in a model system for speciation research. We acknowledge, however, that the evidence for the presence of functional genetic clusters is preliminary. The inference of genetic clusters is limited by a number of methodological constraints. First, our annotations are mostly based on *D. melanogaster* proteins, which are rather divergent from crickets, and hence depend on the presence of conserved regions. We therefore have incomplete identification of homologs and GO annotations. In addition, we can only annotate scaffolds that are in the linkage map, which are, in each case, inherently a subset of the scaffolds that make up a given genomic region. However, it is not apparent to us that the sampling we are able to do, while incomplete, would bias our results in favor of the GO enrichment and gene clustering we observed.

Genomic clustering of causal loci controlling species differences in pulse rate would have profound consequences on the evolution of *Laupala* mating behavior during speciation. Genomic clustering of genes has been associated with several traits that are important in reproductive isolation [28,101], speciation [102], and mating behavior variation [25–27]. Gene clusters would offer a potential adaptation to overcome constraints associated with behavioral evolution, which surely requires coordinated changes in many loci controlling complex neurophysiological traits. Close linkage would reduce interspecific and, potentially, intraspecific recombination and facilitate co-adaptation [23]. For mating behavior, not only linkage of multiple song genes, but also of song and preference genes could speed up divergence and speciation [103,104]. In *Laupala* there is evidence for co-localization of male song and female preference QTL [105,106]. Linkage and orchestrated evolution of different song genes and of song and preference genes might be facilitated by reduced recombination and the co-adaptive gene clusters might contribute to the rapid divergence of mating behavior in the young but diverse radiation of *Laupala*.

### 4.4 Candidate genes

Our results also suggest potential candidate genes that control mating behavior variation in *Laupala*. The main enriched biological processes among the predicted gene products in QTL regions can be tied to potential modulators of central pattern generators (driving rhythmic behaviors) and to sex-specific expression of nervous system development pathways in fruit flies. These findings were in part driven by potential homologs of the motor neuron development gene *roundabout* (1.5 cM away from the peak at LG1), a Leucin-rich repeat kinase involved in neurodegenerative disease and locomotion located 0.5 cM from the peak on LG3, *lola* (transcription factor regulating neuromuscular development 6 cM away from the peak at LG4), and *semaphorin* (directly flanking the peak at LG7). All but the Leucin-rich repeat kinase are affected by sex-specific transcription of *fruitless* in *D. melanogaster* (located on LG2 in *Laupala*) and contribute to sexual dimorphism in the nervous system [107–109].

In addition to these sex specific receptor proteins, we find receptors for serotonin, GABA, dopamine, and acetylcholine, all known neuromodulators of central pattern generators in insects [43,110,111]. We also identify several ion channel genes such as *cadherin* (flanking the QTL peak marker on LG3), *KCNQ potassium channel* (1.5 cM from peak marker on LG3), *cacophony* (16 cM from the peak on LG4, which has a functional role in the inter-pulse interval in *D. melanogaster* [112]), and *sandman* (1.9 cM from the peak at LG5). Without functional evidence, however, we can only consider these genes as candidate loci and cannot speculate further about the genetic and neurobiological pathways involved in song generation and song differentiation in *Laupala*.

## 5. Conclusions

Together, this study presents rare comparative insights into the polygenic genetic architecture associated with sexual trait divergence during speciation in different biogeographic contexts. Clearly, rapid quantitative trait differentiation associated with speciation can occur under a Type I genetic architecture, where many genes diverge in concert to produce a conspicuous species difference. We show that the genetic architecture of male song rhythm divergence in closely related *Laupala* species is remarkably similar among two independently originated species pairs, with comparable QTL numbers, effect sizes and an overall absence of interaction among loci, despite their different geographic histories. These similarities may in part result from constraints on the spatial distribution of genetic variation controlling pulse rate divergence due to clustering of causal loci. We show that the identified QTL regions underlying song pulse rate divergence are enriched for a variety of neuromuscular processes potentially contributing to modulating the central pattern generators that control the song rhythm. This enrichment pattern suggests a compelling genetic potential, deriving from the clustering of multiple, physically linked loci, for rapid divergence in *Laupala* mating behavior. We further identify several potential candidate genes controlling a highly divergent behavioral phenotype that forms a major barrier between recently diverged species.

## Supplementary Materials

The following will be made available online Figure S1: Linkage map and multivariate QTL model results, Figure S2: Inversion on LG1, Table S1: QTL dominance effects, Table S2: QTL peak and flanking markers, Table S3: Annotated transcripts mapping to QTL regions after correcting for pseudo-replication, Table S4: GO enrichment of transcripts in Table S3, Table S5: GO enrichment of transcripts in Table S3 per linkage group, Table S6: Annotated transcripts mapping to QTL regions, Table S7: GO enrichment of transcripts in Table S6, Table S8: GO enrichment of transcripts in Table S6 per linkage group.

## Author Contributions

“Conceptualization, T.B., K.P.O., and K.L.S.; Formal Analysis, T.B.; Investigation, K.P.O. and K.L.S.; Writing-Original Draft Preparation T.B.; Writing-Review & Editing, K.P.O. and K.L.S.; Visualization, T.B. and K.L.S.; Supervision, K.L.S.; Project Administration, K.L.S.; Funding Acquisition, K.L.S.”

## Acknowledgments

We would like to thank Jon Lambert for sample collection and RNA extraction, Michael Brewer for assistance with transcriptome sequencing, and Aureliano Bombarely for help with the bioinformatic analyses that contributed to the transcriptome assembly. We thank the Shaw Lab, and especially Mingzi Xu, for valuable discussions of methodology. Furthermore, we acknowledge Karl Broman and the R/QTL discussion group as well as the Biostars and Stackoverflow communities for discussion that helped implement the analyses.

## Supplementary Material

Fig S1, Fig S2: see below

Table S1-S8, available upon request.

**Figure S1.**
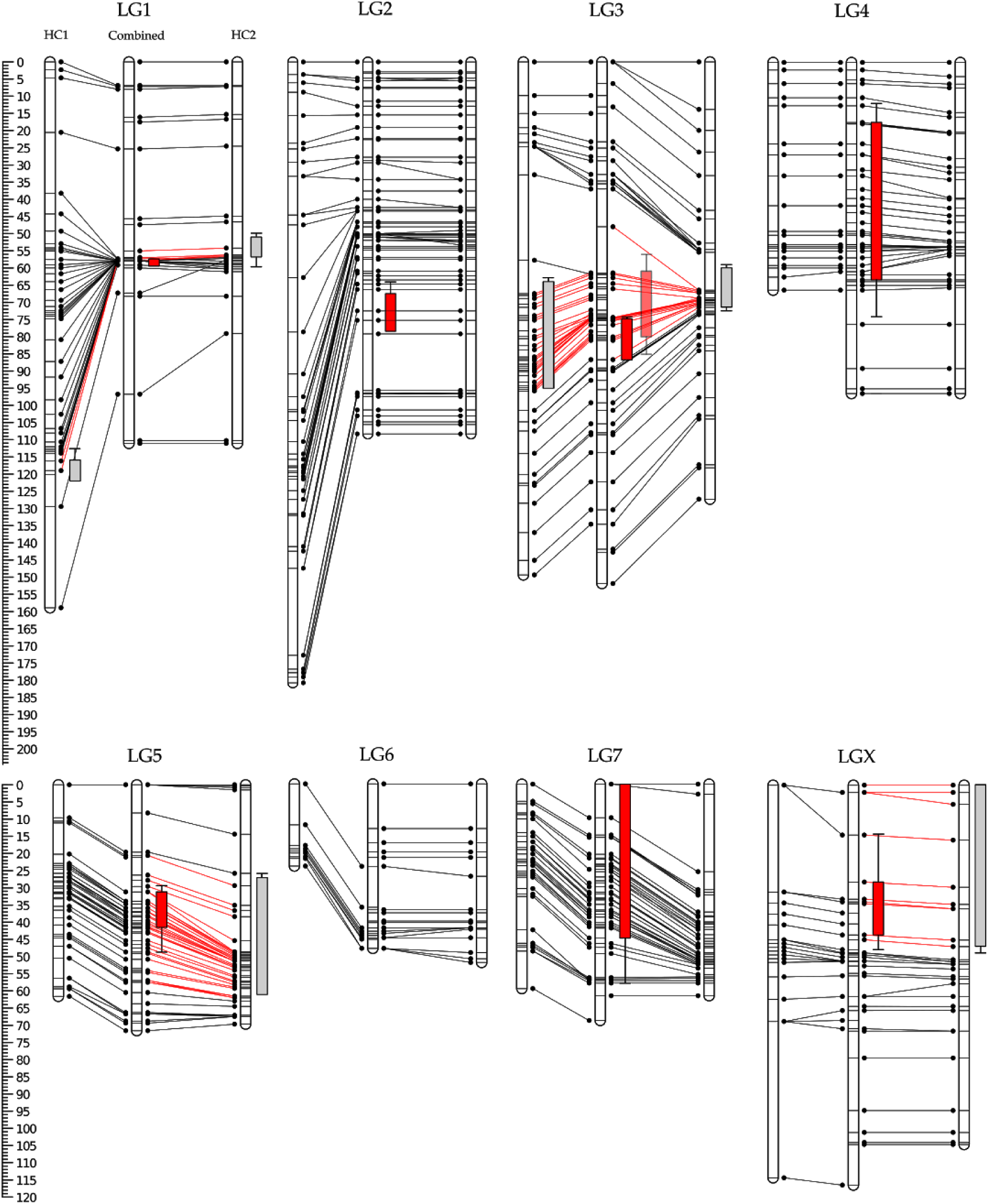
Linkage map and muliivariate QTL model results. The horizontal black bars show the position of the SNP markers in the HC1, combined, and HC2 map, respectively. The black dots and solid lines connect markers on scaffolds shared between adjacent Linkage groups. The scale indicates the position along the linkage groups in cM. The grey/red vertical rectangles and error bars show the 1-LOD and 1.5-LOD interval around the QTL peaks, respectively. We adjusted the 1.5-LOD interval for the HC1 QTL on LG1 to correct for the “phantom” peak at 37 cM. Red lines connect shared scaffolds within the -LOD intervals. On LG3, the partially transparant LOD interval indicates the position of the LOD interval in the model when only considering QTL on LG1, 3, 5, and X.

**Figure S2.**
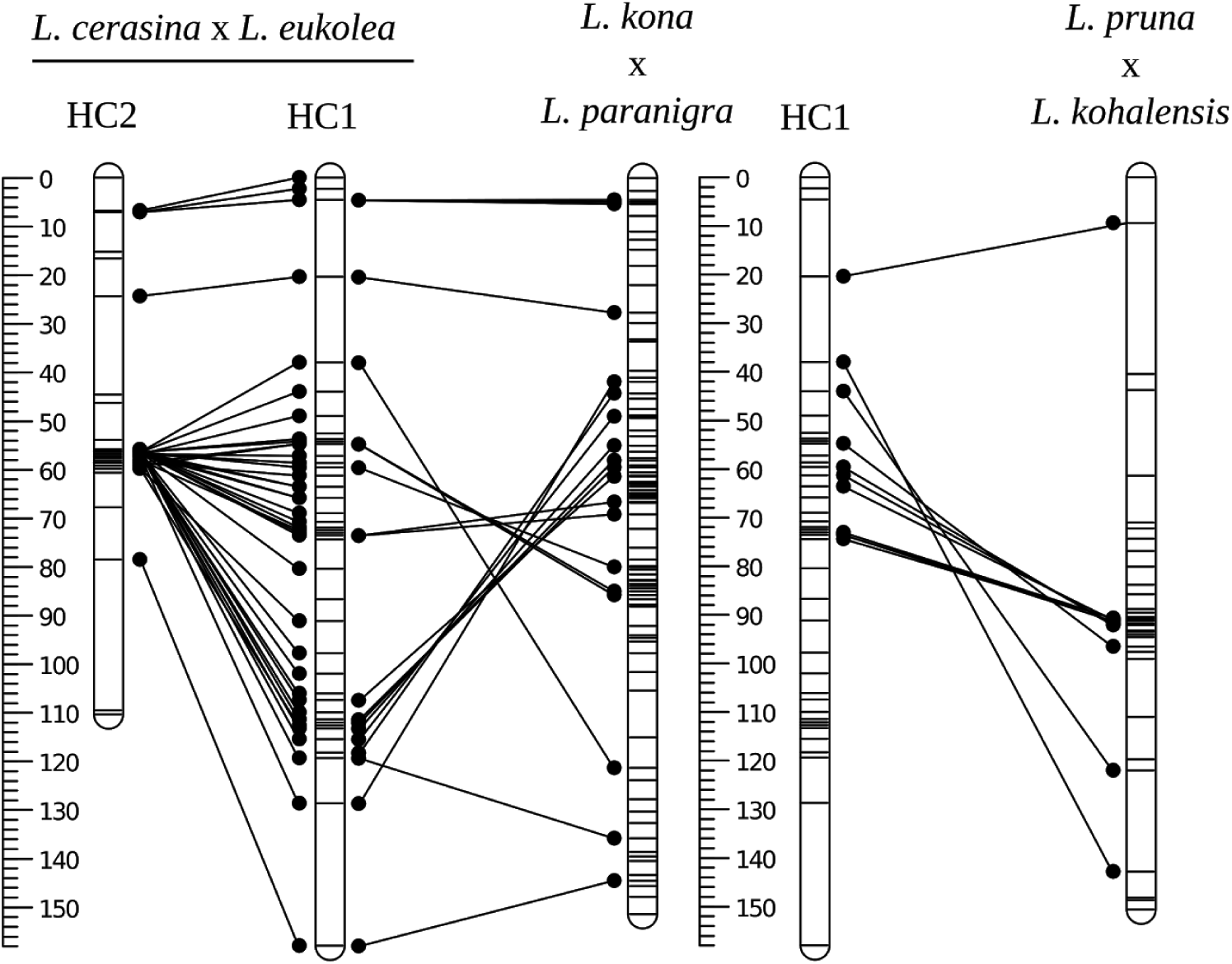
The region of extremenly low recombination in HC2 relative to HC1 appears to be due to crossing parental lines with alternatively oriented inversion karyotypes. Comparing the marker order in HC1 with the marker order on homologous linkage groups in two other interspecies linkage maps (see Blankers et al. 2018 for details), shows that the marker order in HC 1 is inverted relative to the other maps. Although further investigation is needed, the pattern suggests that a large pericentric inversion is segregating in either L. cerasina or L. eukolea, or both.

